# Isolation of a putative S-layer protein from anammox biofilm extracellular matrix using ionic liquid extraction

**DOI:** 10.1101/705566

**Authors:** Lan Li Wong, Gayathri Natarajan, Marissa Boleij, Sara Swi Thi, Fernaldo Richtia Winnerdy, Sudarsan Mugunthan, Yang Lu, Jong-Min Lee, Yuemei Lin, Mark van Loosdrecht, Yingyu Law, Staffan Kjelleberg, Thomas Seviour

## Abstract

Anaerobic ammonium oxidation (anammox) performing bacteria self-assemble into compact biofilms by expressing extracellular polymeric substances (EPS). Anammox EPS are poorly characterized, largely due to their low solubility in typical aqueous solvents. Pronase digestion achieved 19.5 ± 0.9 and 41.4 ± 1.4% (w/w) more solubilization of *Candidatus* Brocadia sinica-enriched anammox granules than DNase and amylase respectively. Nuclear magnetic resonance profiling of the granules confirmed that proteins were dominant. We applied ionic liquid (IL) 1-ethyl-3-methylimidazolium acetate and N,N- dimethylacetamide (EMIM-Ac/DMAc) mixture to extract the major structural proteins. Further treatment by anion exchange chromatography isolated homologous S/T-rich proteins BROSI_A1236 and UZ01_01563, which were major components of the extracted proteins and sequentially highly similar to putative anammox surface-layer (S-layer) protein KUSTD1514. EMIM-Ac/DMAc extraction enriched for these putative S-layer proteins against all other major proteins, along with six monosaccharides (i.e. arabinose, xylose, rhamnose, fucose, galactose and mannose). The sugars, however, contributed <0.5% (w/w) of total granular biomass, and were likely co-enriched as glycoprotein appendages. This study demonstrates that S-layer proteins are major constituents of anammox biofilms and can be isolated from the matrix using an ionic liquid-based solvent.

## Introduction

Anaerobic ammonium oxidation (anammox), whereby ammonium is anaerobically oxidized directly to nitrogen gas with nitrite as electron acceptor, contributes up to 80% of oceanic N losses^1^, and when coupled with partial nitrification is a more sustainable option for N removal from waste waters than conventional processes^2^. Anammox has been researched intensively over several decades across different natural environments as well as more than 100 full-scale wastewater treatment plants^3^. Nineteen anammox species have been identified so far with all belonging to the *Planctomycetes* phylum^4^. Each species, regardless of habitat, self-assembles through extracellular polymeric substances (EPS) expression, into either supported biofilms or granules^5^. Biofilm existence allows anammox bacteria to form syntrophic relationships with other bacteria (e.g. ammonium oxidizing bacteria)^6^ and enhances sludge retention. EPS can also increase microbial tolerances to a range of environmental stresses, such as shear, oxidative and salinity stress^7^.

EPS have key roles in anammox biofilms. Little is known about the structure and function of specific EPS in water and wastewater biofilms^8^. Research into anammox EPS has largely focused on general classes of molecules^9^, for example changes in total protein or sugar content in response to environmental conditions like shear^10^ and salinity stress^11, 12^. *In situ* characterizations have revealed the spatial distribution of proteins, polysaccharides and cells in anammox sludges. For example, loose protein secondary structure in anammox EPS has been shown to expose large amounts of hydrophobic amino acid groups for hydrophobic interactions that effectively mediate anammox EPS aggregation^13^. Similarly, a *β*-sheet structure of extracellular proteins in anammox granules was found to maintain the function of inner hydrophobic groups, while uronic acids further support the biofilm matrix and prevent cell detachment^14^. Nonetheless, the precise identities and compositions of anammox EPS remain undescribed.

Identifying key biopolymers within anammox EPS has always been a major challenge due to their poor solubility in conventional, typically aqueous, extraction solvents. In addition, a variety of anammox biofilm morphologies, including homogeneously-distributed^15, 16^, stratified granules^17–20^, flocs^21^ and surface-attached^22^ structures further confound the extraction and characterization of anammox EPS. Numerous attempts to extract anammox EPS through either cationic exchange resin (CER)^23^, physical methods such as centrifugation, heating and sonication or chemical methods by using organic solvents (detergents and ethanol), have been made. However, to date no exopolymer has been isolated from the matrix of anammox biofilms, which is a minimum requirement for subsequent biophysical and functional analysis.

While some metabolic proteins are conserved between anammox species (e.g. the c-type heme proteins)^24^, it is unknown whether any extracellular proteins are similarly conserved. It has been reported that surface layer (S-layer) proteins may be common to anammox biofilms as a structural component of the cell. For example, KUSTD1514 forms a shell on the outside of the cell envelope of *Ca.* Kuenenia stuttgartsiensis^25^, while a homologous S-layer protein is a major EPS constituent of *Ca.* Brocadia-enriched granular biofilm and hypothesized to also contribute to biofilm matrix structure (i.e. following shedding)^26^. S-layer proteins could be important to biofilm formation more broadly^27^. Describing a function for S-layer proteins in biofilm formation is confounded by the fact that they are embedded in EPS and the S-layer protein has not been isolated from the anammox biofilm matrix.

To address the challenge of processing anammox EPS, we used ionic liquid 1-ethyl-3-methyl imidazolium acetate (EMIM-Ac) to dissolve a laboratory anammox granular biofilm. Imidazolium-based ionic liquids are green solvents that have also been applied to process similarly recalcitrant biopolymers (e.g. cellulose and chitosan)^28^, as protein stabilizers and co-solvents, and enzyme catalysts. We extracted and purified the putative S-layer protein from a laboratory anammox granular biofilm. We found a putative S-layer protein to be the dominant polymer in our biofilm extract. While six monosaccharides were co-enriched with the EPS, they contributed <0.5% (w/w) of total granular biomass and were all common protein o-glycosylating sugars. It is thus likely that they were enriched as glycoprotein appendages rather than free exopolysaccharides. We provide further support for an important role for S-layer proteins in anammox biofilms, and present a method, involving EMIM-Ac, that allows for S-layer proteins to be isolated from complex matrices such as biofilm EPS. This will inform on the role of S-layer proteins in anammox biofilm formation.

## Materials and methods

### Anammox granular sludge enrichment

Anammox granular sludges (GR) were cultivated in a 4 L reactor seeded with activated sludge from a full-scale water reclamation plant (WRP) in Singapore and fed a synthetic medium consisting of (g/L): KHCO3 1.25, KH2PO4 0.025, CaCl2∙6H2O 0.3, MgSO4∙7H2O 0.2 and FeSO4∙7H2O 0.025. More details on laboratory anammox granules enrichment are provided in the Supporting Information (SI).

Granules were extracted from the reactor immediately prior to the settle and decant phase, washed with double-distilled water and lyophilized (FreeZone Plus 4.5 Liter Cascade Benchtop Freeze Dry System).

### Enzymatic treatment of granules

Granules (GR) were digested by Pronase E, DNase A, and α and β amylase (Sigma Aldrich, Singapore), either independently or in sequence. 2 mg/mL of each enzyme was applied to 10 mg/mL biofilm in 2 mL Eppendorf tubes. An incubation time of 2 h was applied for all the treatments at 37°C (for Pronase E and DNase A) and 60°C (for α and β amylase). Pronase E (0.16% (w/w)) was prepared in 0.1 M Tris, 0.5% SDS, pH 7.5, DNase A (1% (w/w)) was prepared in 0.1 M Tris, 25 mM MgCl_2_, 5 mM CaCl_2_, pH 8.0 while α and β amylase (25 μL/mL and 2.5 μL/mL) were prepared in 16 mM sodium acetate buffer, pH 6. After digestion, the supernatant was removed and the residual solids washed three times with double distilled water. Total suspended solids (TSS) and volatile suspended solids (VSS) were measured according to the APHA standard method^29^. TSS and VSS solubilization percentages were calculated as the percentage difference in TSS and VSS respectively before and after enzymatic treatment.

### EPS extraction by ionic liquid

The method for extracellular polymeric substances (EPS) extraction and processing is summarized in Figure 1.

**Figure 1:**
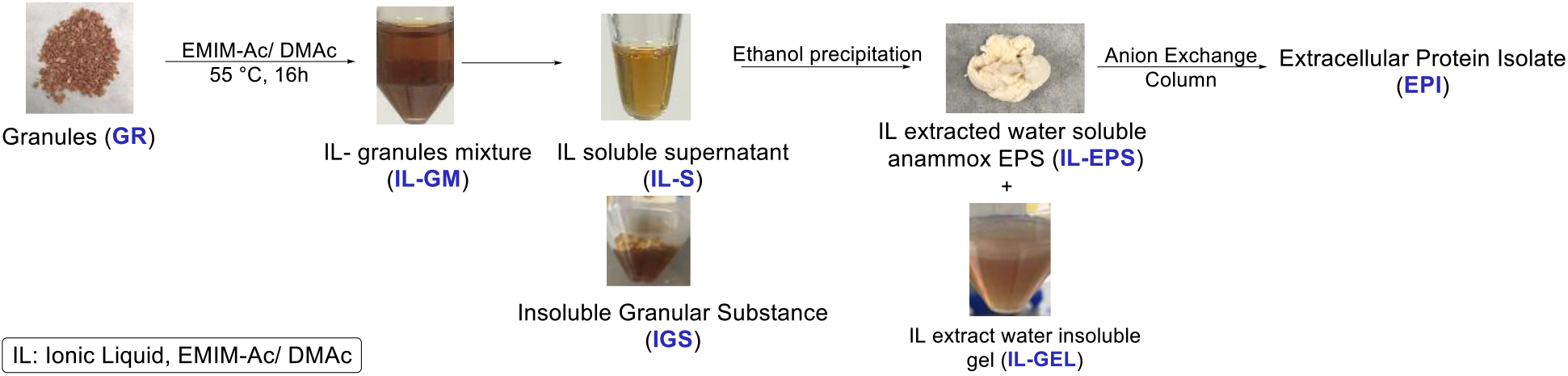
Schematic diagram of anammox extracellular polymeric substances (EPS) extraction and purification with ionic liquid (IL)

Freeze-dried (FreeZone Plus 4.5 Liter Cascade Benchtop Freeze Dry System) laboratory anammox granules (GR) were directly added to 40:60 (v/v) 1-ethyl-3-methyl imidazolium acetate (EMIM-Ac)/N,N-dimethylacetamide (DMAc) mixture as described by Seviour et al.^30^, to a concentration of 30 mg dry sludge/mL. The tube was incubated in a water bath at 55°C for 16 h. The ionic liquid-soluble fraction (IL-S) was captured by precipitation with ethanol (70% (v/v)), separated by centrifugation, cleaned by dialysis and lyophilized to give IL-extracted anammox EPS (IL-EPS) for further analysis.

### Nuclear magnetic resonance (NMR) spectroscopy of acid-digested granules

The IL-EPS fraction (37.5 mg/mL) was dissolved in 4 M trifluoroacetic acid-d (TFA-d) and heated to 120°C for 5 h. 1D ^1^H and 2D ^1^H-^13^C HSQC NMR spectra of acid-digested IL-EPS were then collected on a 700 MHz Bruker Avance II spectrometer at 25°C. Due to the high concentration of IL-EPS precipitate in the 4 M TFA-d, the required pulse length for the 90° excitation pulse was extremely long. Spectra analyses were performed using Topspin 4.0 (Bruker) software.

### ^31^P NMR analysis of soluble EPS

^31^P NMR solution state NMR experiments were performed on a 400 MHz Bruker Avance spectrometer at 25°C. Asolectin (30 mg/mL) (Sigma Aldrich) and anammox granules (30 mg/mL) for NMR analysis were prepared in 40:60 (v/v) EMIM-Ac/DMAc. *Pseudomonas aeruginosa* planktonic cell lysate (30 mg/mL) was prepared by lysing the cells in lysozyme. 10% (v/v) of D_2_O and 7.5 mM of H_3_PO_4_ was added to all samples prior to NMR analysis for locking and referencing purposes. All spectral analyses were performed using Topspin 4.0 (Bruker) software.

Briefly, overnight PAO1 WT planktonic cell pellet was resuspended in 2.25 mL of STET buffer (10 mM Tris-HCl, pH 8 with 0.1 M NaCl, 1 mM EDTA, and 5% (w/v) TRITON^®^ X-100) followed by the addition of 258 μL, 10 mg/mL lysozyme (Sigma-Aldrich) in lysozyme solution (10 mM Tris-HCl, pH 8). The mixture was vortexed and incubated in 37°C for 3 hours followed by probe sonication (SM Vibracell CVX750 Probe Ultrasonicator) at 3s interval for 12 min. The lysate pellet was then freeze dried for NMR analysis.

### Staining and microscopy

Microscopic imaging was conducted on a confocal microscope Leica SP8WLL and Zeiss LSM 780 with a 63× objective. Briefly, crude and IL-treated anammox granules were stained with BacLight Live/Dead viability stain (Thermo Fisher Scientific) according to manufacture’s manual. IL treated anammox granules (IM) were washed twice with double distilled water and freeze-dried before imaging.

### Anion exchange column

IL-EPS was dissolved in 20 mM HEPES, pH 8.0 and passed through 0.2 μm filter prior to being passed through purified anion exchange column connected to Amersham Akta fast protein liquid chromatography (FPLC). The IL-EPS mixture was loaded onto a Source15Q column pre-equilibrated with HEPES and eluted with a linear salt gradient of 95-275 mM NaCl. Fractions containing the targeted protein were combined and concentrated by centrifugation filter, molecular weight cut-off 3 kDa (Amicon®, Merck) to a concentration of 1 mg/mL as determined by Qubit Protein Assay Kit (Invitrogen™) and the Qubit 3.0 Fluorometer (Invitrogen™).

### SDS-PAGE of the soluble EPS

20 μL of the IL-EPS and fractions eluted from the anion exchange column by 245 and 280 mM NaCl (EPI F6, EPI F7) were denatured in Laemmli buffer (Bio-Rad) (1:1 (v/v)) for 10 min at 98°C and were loaded onto pre-cast NuPAGE 4-12% (w/v) Bis-Tris 1.5-mm minigel (Invitrogen™). The electrophoresis was carried out at 180 V in NuPAGE MES SDS running buffer (50 mM Tris base, 50 mM MES, 0.1% (w/v) SDS, 1 mM EDTA. pH 7.3) for 40 min. After electrophoresis the gel was removed from the cassette, washed multiple times with double distilled water and stained with Coomassie Blue (1g Coomassie Brilliant Blue in methanol (50% (v/v)) and glacial acetic acid (10% v/v))) for 1 h. The gel was then destained with destaining solution (25% (v/v) methanol, 5% (v/v) acetic acid) until protein bands became visible. Glycoprotein was detected by periodic acid-Schiff (PAS) staining kit (Thermo Fischer Scientific Pierce Glyocoprotein Staining Kit).

### LC-MS/MS analysis of gel bands

200 and 170 kDa bands were excised from the SDS-PAGE gel and subjected to reduction, alkylation and in-gel tryptic digestion as described by Shevchenko et al.^31^. Following digestion, the peptides were eluted with 1:1 (v/v) water/acetonitrile (ACN) solution with 0.2% (v/v) TFA into a microtiter receiver plate by vacuum and then concentrated by vacuum centrifugation (miVac, SP Scientific).

Peptide analysis was performed by LCMS-TripleTOF 5600 (refer to Supporting Information for details). Peptides were identified using ProteinPilot 5.0 software Revision 4769 (AB SCIEX) with the Paragon database search algorithm (5.0.0.0.4767) and the integrated false discovery rate (FDR) analysis function. These protein data were searched against a protein reference database of translated predicted genes from metagenome assemblies constructed from previously sampled reactor gDNA. The metagenome contained a recovered complete genome of *Ca.* Brocadia^32^ and also included five extant draft AnAOB genomes, enabling comparative analysis.

### Antibody generation

Two rabbits were inoculated with polypeptide sequence cys-DIREITGVASDR, representing amino acid sequence 742-753 of BROSI_A1236 and identified to be an exposed region of the protein^33^, in a three-month immunization protocol consisting of four injections on days 0, 30, 50 and 80 (SABio, Singapore). The serum was collected and affinity purified from rabbit antiserum. Total IgG fractions (crude serum) of one rabbit were used as primary anti-BROSI_A1236 antibody. Refer to Figure S1 for Dot Blot analysis of the rabbit sera binding to the BROSI_A1236 polypeptide.

### Immunoblot analysis

Proteins were transferred from the SDS-PAGE gel to a membrane using iBlot transfer system (Invitrogen™). The polyvinylidene difluoride (PVDF) membrane was blocked with PBS-T (137 mM NaCl, 12 mM PO_4_^3−^, 2.7 mM KCl, 0.05% Tween®20, pH 7.4) and 5% (w/v) bovine serum albumin (BSA) for 1 h at 22°C. The primary antibody was then diluted 3000 times in blocking buffer and incubated for 2 h at 22°C. PVDF membrane was washed three times with PBS-T for 10 min before incubating with Goat anti-Rabbit IgG (H+L) Secondary Antibody, HRP (ThermoFisher Scientific) diluted 5000 times in blocking buffer for 1 h at 22°C in the dark. After incubation, the membrane was washed five times for 15 min with PBS-T. For immune detection, the blot was then developed in 1:1 (v/v) dilution of SuperSignal™ West Femto Trial Kit (ThermoFisher Scientific) to achieve the desired signal intensity in the Amersham Hypercassette™ Autoradiography Cassettes.

X-ray film (CARESTREAM Medical X-ray Green/MXG Film) was exposed to the blot in the cassette from 2 s to 10 min, depending on the intensity of the signal. The film was then inserted into an auto processor provided by Abnova for film development.

### Size exclusion chromatography (SEC) analysis with fluorescence detector

SEC was performed using (i) a liquid chromatograph with pump (LC-20AD), (ii) a fluorescence detector (RF-20AXS), (iii) an auto sampler (SIL-20AHT), and (iv) a communication module (CMB-20A). The molecular weights (MW) of the EPS samples were estimated by passing 15 μL of the filtered samples through an analytical scale SEC column (OHpak SB-804 HQ) following its guard column (OHpak SB-G). Tris buffer (25 mM Tris, pH 7.0 ± 0.1) was used as the mobile phase. The SEC column was calibrated using transferrin, serum albumin bovine, myoglobin and beta amylase obtained from Sigma Aldrich (the details are provided in Supporting Information (SI)).

### Global protein analysis of *Ca.* B. sinica-enriched granules

Protein extraction was performed as previously described^34^. Briefly, lyophilized anammox granules were resuspended in extraction buffer and cells were lysed by bead beading. Cell debris was removed, and extracted proteins were precipitated overnight using trichloroacetic acid (TCA). Pellets were washed three times in 100% ice-cold acetone, dried, and resuspended in 100 mM HEPES (pH 7.5). Protein concentration for each sample was estimated in a single technical replicate using the Qubit Protein Assay Kit (Invitrogen) and the Qubit 3.0 Fluorometer (Invitrogen).

Protein samples were then subjected to preparative denaturing SDS-PAGE (refer to Supporting Information). Gels were Coomassie-stained (Bio-Safe Coomassie Stain, Bio-Rad) and the protein bands (1 × 1 × 0.08 cm) excised and subjected to reduction, alkylation and in-gel tryptic digestion, as described elsewhere^31^. Following digestion, peptides were recovered, dried and reconstituted in 5% formic acid prior to desalting and clean-up using StageTips^35, 36^ prepared as previously described^37^ with the modification that an additional layer of PorosOligo R2 material (Applied Biosystems, Foster City, CA, USA) was added on top of the PorosOligo R3 material. Following purification, peptides were eluted using 66% (v/v) ACN, dried using vacuum centrifugation and reconstituted in 0.1% (v/v) TFA/2% (v/v) ACN solution.

Peptides were subsequently analyzed by ultra-high-performance liquid chromatography (UHPLC) using an Easy-nLC1200 (Thermo Scientific) online system coupled to a Q Exactive High Field mass spectrometer (Thermo Scientific) (refer to Supporting Information for complete details). Raw data from the Q Exactive High Field were analyzed using MaxQuant v. 1.6.3.4. Each experiment was defined as its own parameter group. The label-free quantification (LFQ) and the intensity based absolute quantification (iBAQ) features were enabled, and LFQ was separated in parameter groups. Oxidation of methionine was set as a variable modification, and carbamidomethylation of cysteine was set as a fixed modification. The ‘match between runs’ feature was selected, while all other settings were left as default.

The raw data were searched against an inhouse-generated database containing the open reading frames from *Ca*. Brocadia bins recovered from metagenomics assembly of the *Ca*. Brocadia sinica-enriched reactor^32^.

### Carbohydrate analysis

Carbohydrate analysis of *Ca.* B. sinica granules (GR) and the IL-EPS samples extracted from *Ca.* B. sinica granules, was performed by Shimadzu Gas Chromatography triple quadrupole mass spectrometer (GC-QqQ-MS/MS) in multiple reaction monitoring (MRM) mode. The samples were digested in 4 M TFA for 4 h at 100°C^38^.

## Results

### Profiling the contribution of EPS classes to anammox biofilm structure

*Ca.* B. sinica-enriched granules (GR) (Figure S2) were digested with enzymes targeting general classes of extracellular polymers commonly found in environmental biofilms, including proteins (P), DNA (D) and α- and β-glycans (A) (Figure 2A). Sludge from the laboratory-scale reactor enriched with *Ca.* B. sinica had a higher VSS content of 88% compared to industrial granules^39^. All enzymes achieved a higher degree of solubilization of the organic fraction relative to total biofilm, as indicated by low and high reductions in TSS and VSS respectively, following enzymatic digestions. This is likely due to the presence of inorganic minerals, which are common features of anammox biofilms (e.g. apatite)^40^. Digestion with pronase achieved a higher degree of solubilization than either DNase A, α and β amylase or the control (i.e. heating only at 60 °C). A similar amount of VSS and TSS solubilization was observed in the control sample as for α- and β- amylase-treated granules (i.e. same buffers and heated to the same temperature). Thus, neither α- or β-sugars contributes to the structure of the anammox EPS. Furthermore, high EPS solubilization is consistently achieved for granules treated with pronase, either by itself or coupled with DNase A, α- or β-amylase. This suggests that proteins are important structural components of the *Ca.* B. sinica-enriched anammox EPS matrix.

**Figure 2:**
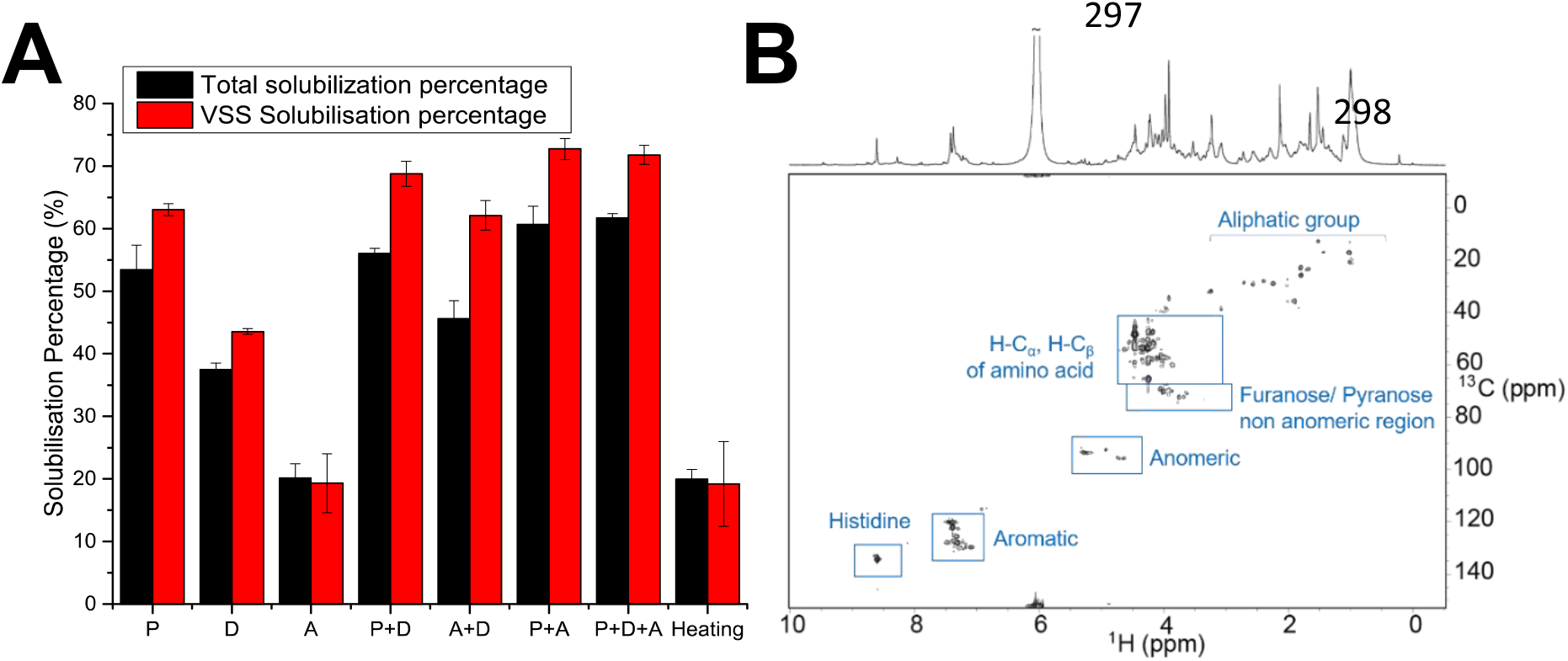
(A) Percent solubilization of volatile and total solid in *Ca.* B. sinica-enriched granules (GR) grown in a laboratory reactor following digestion with enzymes: pronase E (P), DNase A (D), α or β amylase (A). (B) ^1^H-^13^C HSQC and 1-D ^1^H NMR spectra of acid-digested anammox granules EPS (IL-EPS) in 4 M TFA-d (37.5 mg/mL) at 25°C showing a distribution of ^1^H-^13^C HSQC cross peaks that is consistent with the presence of protein and hexose-based sugars.

The heteronuclear single quantum coherence (HSQC) NMR spectrum of acid-digested anammox granule EPS (IL-EPS) (Figure 2B) shows that some polysaccharides were present along with proteins. This is illustrated by HSQC cross peaks at (δ_H_, δ_C_) of (4.8-5.3, 92.4-95.5), consistent with the anomeric region of polysaccharides^41^ (δ_H_ and δ_C_ are the ^1^H and ^13^C chemical shifts in ppm respectively). It is therefore possible that polysaccharides not targeted by α- or β- amylase co-exist with structural proteins (i.e. not α- or β-glycans such as granulan or alginate-like exopolysaccharides)^42^. Another explanation could be that glycoproteins are a component of the anammox biofilm EPS. Nonetheless, the spectrum is dominated by HSQC cross peaks at (δ_H_, δ_C_) of (4.2-4.8, 45.6-67.5), (7.2-7.6, 119.1-131.7) and (8.5-8.7, 130.7-136.7), which represent amino acid α-regions and aromatics, and histidines, respectively, further supporting our finding that proteins dominate *Ca.* B. sinica-enriched granules^43, 44^.

### Ionic liquid-based EPS extraction

We therefore aimed to extract structural proteins from the *Ca.* B. sinica-enriched granules, and used ionic liquid 1-ethyl-3-methyl imidazolium acetate (EMIM-Ac) for this purpose given its demonstrated ability to dissolve recalcitrant polymers. The recovery yields of representative biofilm exopolymers (i.e. basic and acidic proteins, anionic, cationic and neutral polysaccharides) following dissolution in 40:60 (v/v) EMIM-Ac/DMAc with recovery by ethanol precipitation (i.e. anti-solvent), were determined (Figure S3). Recovery yields of 54.1 ± 9.0 and 23.2 ± 8.7% (w/w) for basic and acidic proteins (i.e. cytochrome C and lipase) respectively were achieved following EMIM-Ac/DMAc solubilization and ethanol precipitation. EMIM-Ac/DMAc dissolution with ethanol precipitation is therefore a viable strategy for recovering extracellular proteins from anammox biofilms. Further, EMIM-Ac/DMAc coupled with ethanol as anti-solvent recovered neutral and cationic polysaccharides (i.e. cellulose and chitosan), consistent with what has been described in the literature^45^.

Anammox granules dissolved in EMIM-Ac/DMAc at 55°C (16 h) separated into two phases; an insoluble granular phase (IGS), and an ionic liquid soluble phase (IL-S) (Figure 1). The recovery yield following EMIM-Ac/DMAc solubilization and ethanol precipitation and dialysis was 8.2 ± 2.0 % (w/w). The recovered fraction subsequently separated during dialysis into a water soluble fraction (IL-EPS) and a water insoluble gel (IL-GEL).

### Ionic liquid treatment increases the permeability of, but does not fluidize, *Ca.* B. sinica cells

Ionic liquids are known to kill cells, which is believed to be due to osmotic shocking of cell membranes^46^. Live-dead stain of the anammox biofilms indicated both live and dead cells, even in the non-treated component (Figure 3A). Relative to the control, EMIM-Ac/DMAc treatment disrupted aggregate organization such that clusters of dead cells dominated and were less evenly distributed (Figure 3B). Nonetheless, there were also live and dead cells and the loss of order in how the cells are arranged could indicate that the EPS has been extracted.

**Figure 3:**
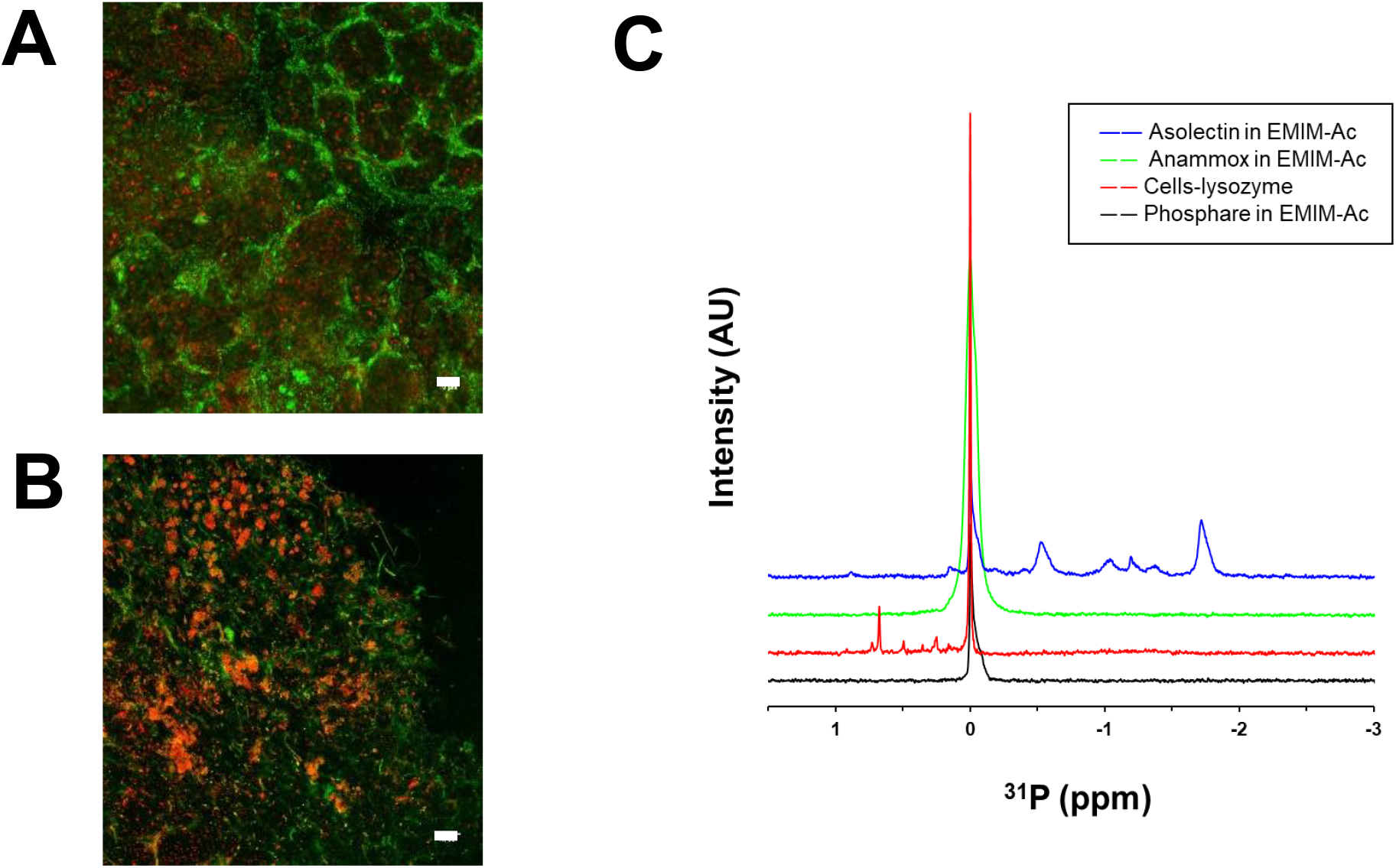
Live (green)-dead (red) stain of *Ca.* B. sinica cells (GR) (A) without any treatment and (B) following dissolution in 40:60 (v/v) EMIM-Ac/DMAc (IM) at 55°C for 6 h. Scale bar indicates 10 μm. (C) ^31^P NMR spectra of azolectin following dissolution in EMIM-Ac/DMAc (55°C, 6 h) (blue), *Ca.* B. sinica-enriched granules following dissolution in EMIM-Ac/DMAc (IS) (55°C, 6 h) (green), lysozyme-treated *Pseudomonas aeruginosa* planktonic cells (red) i.e. a positive control. 7.5 mM phosphoric acid was added to all samples as a reference.

*Planctomycetes* are Gram-negative bacteria with an outer membrane that contains eukaryotic-like phospholipids in the inner leaflet^47^. To investigate whether increased permeability was due to membrane fluidization, we used ^31^P NMR to determine if anammox phospholipids are released into EMIM-Ac/DMAc upon dissolution. The phospholipid standard (i.e. asolectin from soybean) displayed a range of sharp spectral peaks at *δ_P_* −3 − 0 ppm, consistent with diesterified phosphates (Figure 3C, blue), demonstrating that phospholipids dissolved in EMIM-Ac/DMAc can be detected by ^31^P NMR. In contrast, no ^31^P NMR peaks were observed in the spectrum of EMIM-Ac/DMAc following anammox biofilm dissolution (IL-S) (Figure 3C, green) apart from the reference phosphate peak (i.e. H_3_PO_4_) seen at *δ_P_* 0 ppm. To support the hypothesis that cell lysis can be detected by ^31^P NMR, the spectrum of planktonic *Pseudomonas aeruginosa* cells treated with lysozyme showed several sharp spectral peaks in the monoesterified and disesterified phosphate region (Figure 3C, red). The signal observed at *δ_P_* 0 ppm in every sample arose from the 7.5 mM H_3_PO_4_ standard, where the neat H_3_PO_4_ ^31^P NMR spectra is shown in black.

It is possible that the constituents extracted by ionic liquid derived from inside the cells. While the red color of the gel phase following EMIM-Ac/DMAc treatment of the anammox biofilm (Figure 1) could derive from intracellular Cyt-C heme proteins in anammox granules^24^, the ^31^P NMR spectrum nonetheless indicates that the cell membranes remain intact.

### Isolation of extracellular protein

The molecular weight (MW) distribution of anammox biofilm extracellular proteins, recovered from EMIM-Ac/DMAc by ethanol precipitation, was described by SDS-PAGE with Coomassie Blue, periodic acid-Schiff staining (Figure 4A). Major protein bands in the anammox biofilm EPS extract (IL-EPS) (Figure 4A Lane 2) appear at 55, 60, 65, 170 and 200 kDa. A similar protein MW profile was observed in the SDS-PAGE gel of the IL-GEL sample (Figure S4). Thus the same high MW protein doublet dominates both the IL-EPS and IL-GEL, suggesting that it can undergo a sol-gel transition. In subsequent processing of the IL-EPS by anion exchange chromatography (AEC) the high MW doublet of proteins were concentrated in fractions eluted with 0.2M HEPES buffer with NaCl concentrate in the range of 245 and 280 mM (i.e. 170 and 200 kDa, Figure 4A Lane 3). The AEC purified high MW proteins (EPI) also stained positive for periodic acid-Schiff (PAS) stain (Figure 4A Lane 4), indicating that the high MW protein doublet is glycosylated (i.e. glycoproteins)^48^. Gel permeation chromatography further confirmed the purity of the AEC purified protein where the sample was separated into two major peaks across the high MW range (… EPI F6, Figure 4B).

**Figure 4:**
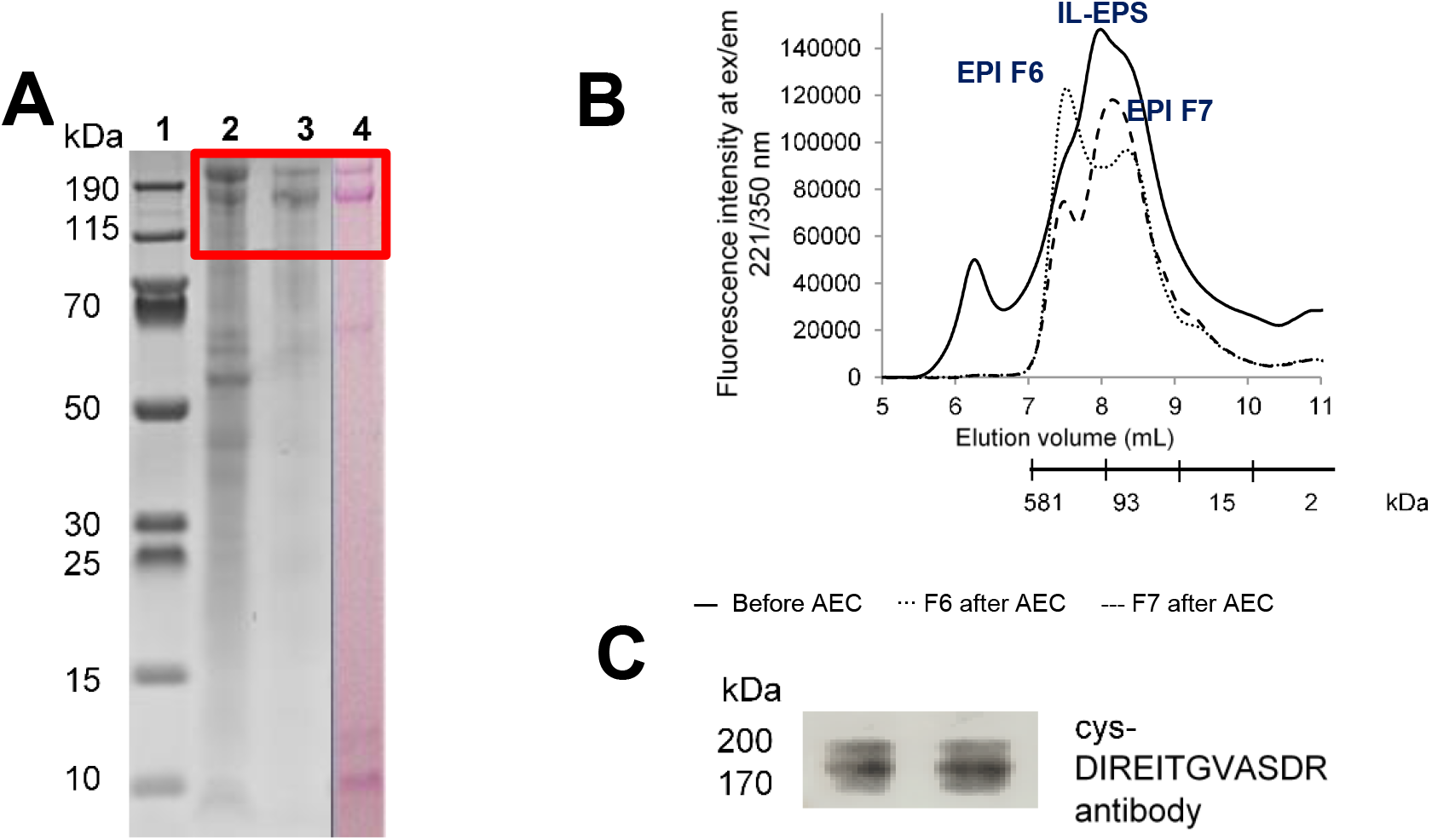
(A) SDS-PAGE gel showing extracellular proteins extracted from *Ca.* B. sinica-enriched granules using EMIM-Ac/DMAc (IL-EPS, Lane 2), anion exchange chromatography (AEC)-purified anammox protein extract (EPI, Lane 3), and periodic acid-Schiff (PAS) stained SDS-PAGE gel showing positively stained high molecular weight glycoprotein doublet (Lane 4). Lane 1 is the PageRuler™ prestained protein ladder. (B) Size exclusion chromatogram profile of crude annamox EPS extract (IL-EPS) and anion exchange chromatography purified EPS fractions (EPI), showing effective AEC purification as well as simplification of chromatogram of sample F6 and F7 after AEC. (C) Immunoblotting validation by Western blot showing positive blot of BROSI_A1236 doublet (EPI) to cys-DIREITGVASDR antibody.

### Co-enrichment of surface layer (S-layer) glycoprotein with commonly o-glycosylating monosaccharides

The AEC-purified proteins (EPI) extracted from *Ca.* B. sinica-enriched granules by EMIM-Ac/DMAc were analyzed by MS/MS and the identified polypeptide fragments mapped against a database for all bacteria. Two hypothetical proteins dominated, BROSI_A1236 and UZ01_01563, that were identical with the exception that BROSI_A1236 has an additional 247 amino acids at the N-terminus^49^. We henceforth refer to them as BROSI_A1236. Immunoblot analysis of the isolated protein, using antibodies raised against amino acids 742-753 of BROSI_A1236, further validated its identity as BROSI_A1236 (Figure 4C; Figure 1). It is highly similar to putative S-layer protein KUSTD1514 (e-value: 0, protein 44%)^50^. The most highly similar abundant proteins in terms of iBAQ (i.e. > 5% abundance) in *Ca.* B. sinica-enriched granules (GR) and the EMIM-Ac extract (IL-EPS) are presented in Figure 5A, with the BROSI_A1236 being the most abundant protein in the EPS prior to AEC purification (i.e. extracted directly by ionic liquid and recovered by ethanol, IL-EPS). There was a two-fold enrichment of BROSI_A1236 following extraction by EMIM-Ac relative to the crude granules, to almost 40% abundance. It is likely, however, that the true abundance of BROSI_A1236 following EMIM-Ac/DMAc extraction was even higher given that mass spectrometry underestimates glycoprotein content due to inhibition of trypsin digestion by the sugars^51^. Hence, EMIM-Ac/DMAc extraction followed by ethanol recovery leads to a high enrichment of a putative S-layer glycoprotein from the extracellular matrix of the anammox granules.

**Figure 5:**
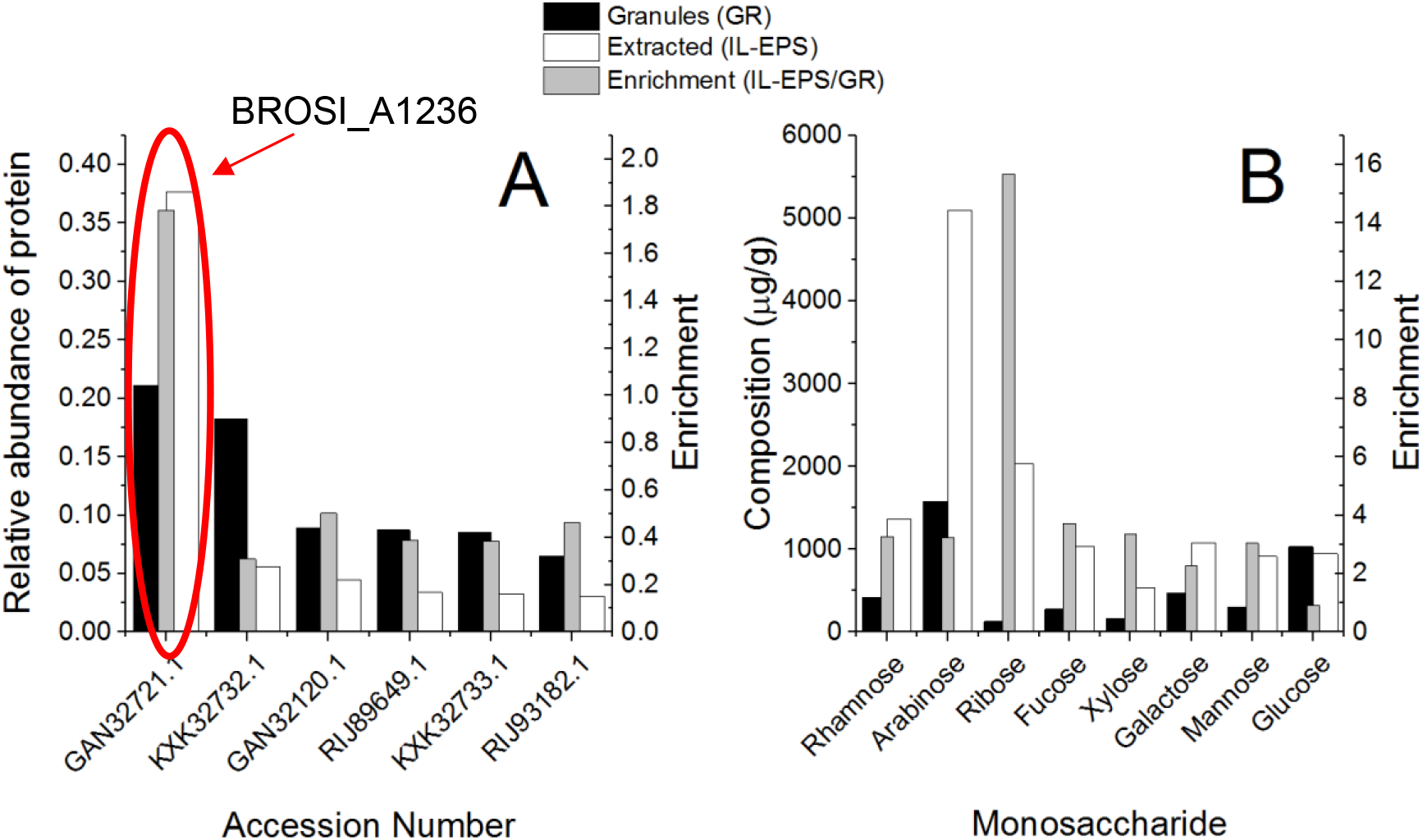
(A) Relative abundance of dominant proteins (i.e. > 5% abundance (iBAQ)) in *Ca.* B. sinica - enriched anammox granules (GR, black) and the water-soluble fraction extracted by EMIM-Ac/DMAc (IL-EPS, white). Degree of enrichment of top six proteins before and after extraction from *Ca.* B. sinica-enriched granules following dissolution in EMIM-Ac/DMAc (grey) (i.e. (iBAQ of water-soluble fraction (IL-EPS))/(iBAQ of granules (GR))). (B) Dry weight composition of typical glycoprotein monosaccharides before and after extraction from *Ca.* B. sinica-enriched granules by EMIM-Ac/DMAc (i.e. composition (μg/g) of water-soluble fraction (IL-EPS)/ composition (μg/g) of granules (GR)). Degree of enrichment of monosaccharides before and after extraction from *Ca.* B. sinica-enriched granules following dissolution in EMIM-Ac/DMAc (grey).

The anammox biofilm contains sugars, as demonstrated by the HSQC NMR spectrum of acid-digested IL extracted anammox EPS (IL-EPS) (Figure 2B). The concentrations of monosaccharides commonly associated with either exopolysaccharides or glycoproteins were measured, and the eight most abundant monosaccharides present in crude anammox biofilms (GR) and in the EMIM-Ac/DMAc soluble material (IL-EPS) are presented in Figure 5B. The fraction of all sugars in the anammox biofilm was 0.45% (w/w), increasing to 1.3% (w/w) in the EMIM-Ac/DMAc extract (IL-EPS). Thus, sugars constitute only a small fraction of the dry weight of the *Ca.* B. sinica-enriched granule (GR), providing further evidence of the importance of proteins in the granule structure. An approximately three-fold increase was observed for all sugars except glucose and ribose, concomitant with a two-fold increase in glycoprotein BROSI_A1236 (EPI). BROSI_A1236 (EPI) is enriched in serine and threonine (11.8 and 16.0% respectively), which are the most commonly o-glycosylated^52^ amino acids. Given that arabinose, xylose, fucose, rhamnose, mannose and galactose are co-enriched with BROSI_A1236 (IL-EPS) in the extracted protein versus the crude anammox biofilm (GR), it is probable that they are appended to the putative S-layer glycoprotein BROSI_A1236.

Two protein bands and GPC peaks were observed from the SDS-PAGE gel and gel permeation chromatogram respectively of the AEC purified fractions (EPI F6 & F7). The high MW peak dominated in EPI F6 from AEC and the low MW peak dominated in EPI F7. These confirm the existence of two structurally similar glycoproteins (i.e. BROSI_A1236 and UZ01_01563), or reflect the different proteoforms of a single protein (e.g. in different glycosylation states).

## Discussion

We observed that proteins are key structural components of *Ca.* B. sinica-enriched biofilms and demonstrated a means to extract and concentrate a putative S-layer glycoprotein using ionic liquid 1-ethyl-3-methyl-imidazolium acetate (EMIM-Ac), which allowed for its subsequent isolation and identification. It is very similar to putative surface layer proteins KUESTD1514 from *Ca.* K. stuttgartsiensis anammox sludge and also to WP_070066019.1 from *Ca.* Brocadia sapporiensis sludges^53^. Furthermore, while sugars were only minor components of the *Ca.* B. sinica-enriched biofilm, they were co-enriched along with the glycoprotein and thus likely exist in the biofilm predominately attached to proteins rather than as free polysaccharides. While S-layers have been postulated to promote biofilm formation, as for *Tannerella forsythia*, the mechanism by which they achieve this is unknown. S-layers may appear as crystalline structures in the matrix as a result of cell-surface shedding^26^. However, it is not clear how crystalline structures might support biofilm formation. The means to extract and subsequently S-layer glycoproteins from biofilm matrices, as described here, will enable methods to be applied to describe the mechanism by which they contribute to the growth of other biofilms. Interestingly, part of the ethanol recovered IL-soluble fraction (IL-S) also formed a gel (IL-GEL) during dialysis. It is therefore possible that, in addition to performing role as a S-layer protein, Brosi_A1236 could also contribute to biofilm formation by undergoing phase separation to a gel. One possible mechanism by which the putative S-layer protein contributes to anammox biofilm could therefore be that it mediates phase separation of the matrix into gels. This could further inform the role of S-layer protein in anammox biofilms, but possibly in biofilms more broadly.

To date, the only reported method of S-layer protein extraction from anammox biofilms was physical extraction by means of Potter homogenizer, which is known to disrupt cells^25^. Chaotropes such as lithium chloride (LiCl)^54^ or guanidine hydrochloride (GnHCl)^55^ are typically used in S-layer protein extractions. LiCl is believed to solubilize S-layer proteins in gram positive bacteria by disrupting the hydrogen bond between protein and secondary cell wall polysaccharides. Such treatments have not been demonstrated for biofilm and EPS solubilization and hence are not likely to be effective at facilitating the isolation of S-layer proteins from extracellular matrices. Ionic liquids can isolate S-layer proteins from biofilm matrices because they are equally effective at solubilizing proteins as they are for recalcitrant polysaccharides like cellulose. Furthermore, they achieve this by disrupting intermolecular hydrogen bonds and increasing solvent order without destabilizing the solute or protein. Similar to LiCl and GnHCl, interactions between the cation and anion of ionic liquids (like EMIM-Ac) have chaotropic effects on the protein and kosmotropic effect on the solvent^56^. The exception to this is cytochrome c, where Fujita et al.^57^ found that ionic liquid had a kosmotropic effect leading to its solubilization. Nonetheless, ionic liquids are good options for isolating extracellular proteins where it is important to preserve their tertiary structure, as would be required to describe the mechanism of the role of S-layer protein in biofilm formation, i.e. to describe their biophysical properties and roles in biofilms, higher order structures and how they interact with cells, other exopolymers and their environment. In addition, ionic liquid EPS extraction was also found here to cause minimal cell disruption as shown by the absence of the phosphate signal in the ^31^P NMR analysis of the ionic liquid EPS extract supernatant (IL-S).

Similarly, the means to solubilize EPS enables a range of approaches to be applied for assigning function, not least of all accurate quantification, but also immunofluorescence microscopy^58^, single molecular structural (e.g. circular dichroism)^59^ and biophysical (e.g. optical tweezing)^60^ analyses. Global quantitative profiling of anammox EPS components has also been applied widely to correlate EPS with certain process parameters, such as salinity resistance^61^, settling performance^62^ and preservation techniques^63^. However, most studies assume that the key EPS are solubilized, which is challenging for the notoriously recalcitrant anammox EPS. Thus, isolating, non-destructively, extracellular polymers from the biofilm will lead to an improved understanding of their function in the anammox biofilm.

We illustrate with this study a non-destructive means to extract extracellular polymers from anammox granules. Additionally, this allows for the subsequent isolation of an S-layer glycoprotein, by anion exchange chromatography, which will enable more detailed structural and functional characterization of a putative S-layer protein from a complex, ecologically and industrially important biofilm.

## Acknowledgements

This research was supported by the Singapore National Research Foundation under its Environment & Water Research Programme and administered by PUB, project number 1301-IRIS-59. SCELSE is funded by Singapore’s Ministry of Education, National Research Federation, Nanyang Technological University (NTU), and National University of Singapore (NUS) and hosted by NTU in partnership with NUS. The authors are thankful to Dr Sharon Longford for proofreading the paper, Dr Henrik Kjeldal and Dr Mads Toft Søndergaard for global protein analysis and Mr Lim Teck Kwang for LC-MS/MS analysis of gel bands.

