## Supplementary material for "Isolation of a putative S-layer protein from anammox biofilm extracellular matrix using ionic liquid extraction": Figure S1

**Materials and Methods**

**Anammox granular sludges enrichment**

Ammonium and nitrite concentrations were increased gradually from 20 mgN/L to 280 mgN/L and 350 mgN/L, respectively over a period of 100 days, with hydraulic retention time reduced gradually from 24 to 16 h in accordance with the nitrogen removal capacity. 1.25 ml/L of trace mineral solution was supplied as described by van de Graaf et al.^1^. Argon/CO_2_ (95/5%) was provided at a constant rate of 25 ml/min during the anoxic phase. Once the nitrogen removal capacity had stabilised the reactor was operated with 12 h cycles comprising of 5 min of feeding, 108 min of anoxic cycle, 67 min of settling and decanting. pH varied between 7.2 to 7.8.

**Size exclusion chromatography (SEC) analysis with fluorescence detector**

Calibration of SEC column was performed according to the method described by^2, 3^. Protein standards with different MW were injected, including transferrin (Sigma Aldrich) – 80 kDa, serum albumin bovine (Sigma Aldrich) – 68 kDa, myoglobin (Sigma Aldrich) – 17 kDa and beta amylase (Sigma Aldrich) – 200 kDa. The logarithm of the MW (in Dalton) shows a linear relationship with the elution volume. The plotted curve for the analytical scale column was regressed to the following equation:

Log aMW = -0,7964Ve + 11,339 (R^2^ = 0,9888)

The plotted curve for the prep-scale column was regressed to the following equation:

Log aMW = -0,0757Ve + 8,6314 (R^2^ = 0,9731)

where aMW is the apparent MW (kDa) and Ve is the elution volume (mL).

**LC-MS/MS analysis of gel bands**

Peptide separation was by Eksigent nanoLC Ultra and ChiPLC-nanoflex columns (Eksigent, Dublin, CA, USA). Samples were desalted (Sep-Pak tC 18 μ Elution Plate, Waters, Miltford, MA, USA) and reconstituted with 20 μl of 98% Water, 2% Acetonitrile, 0.05% Formic acid. They were eluted in an analytical 75 μm×150mm ChromXP C18-CL, 3 μm column (Eksigent, Germany), with peptides separated by a gradient formed by 2% ACN, 0.1% FA and 98% ACN, 0.1% FA. MS analysis was performed on a TripleTOF 5600 system (AB SCIEX, Foster City, CA, USA) in Information Dependent Mode. MS spectra were acquired across the mass range of 400–1250 m/z in high resolution mode (>30000) using 250 ms accumulation time per spectrum. Tandem mass spectra were recorded in high sensitivity mode (resolution >15000) with rolling collision energy on adjustment. Survey- IDA Experiment, with charge state 2 to 4 which exceeds 125 cps was selected.

**Global protein analysis of *‘Candidatus* Brocadia sinica’-enriched granules**

Peptides were first concentrated on a trapping column (Pepmap100, C18, 100 µm × 2 cm, 5 µm, Thermo Scientific), followed by separation on an analytical column (PepmapRSLC, C18, 75 µm i.d. × 50 cm, 100 Å, ThermoFisher Scientific). The temperature of both the trapping column and the analytical column was maintained at 40°C by a butterfly portfolio heater (Phoenix S&T). The Easy nLC1200 was coupled to the QE HF via the nanospray flex source (Thermo Scientific) using distal coated emitter.

The LC gradient used a binary buffer system comprised of 0.1 % formic acid in water (buffer A) and 0.1% formic acid, 80% ACN (buffer b). Peptides were eluted at a constant flow rate of 300 nL/min using the following 120 min gradient: maintaining 5% of buffer b for 1 min, followed by a 3 min ramp to 12 % buffer b, a 90 min ramp from 12% to 37% buffer b, 5 min ramp from 37% to 50% buffer b, 1 min ramp from 50% to 100% buffer maintaining 100 % b for 15 min, decreasing buffer b to 5% over 1 min and keeping that concentration for 4 min.

The mass spectrometer was operated in positive mode, using a data independent top 20 acquisition method. For full MS scan a range 400-1200 m/z was used, setting a resolution of 60000, and an acquired gain control (AGC) target of 1e6 ions. Maximum injection time was set to 50 ms. For fragment scans, the isolation widows was set to 1.2 m/z, resolution to 15000, AGC target was set to 1e5 ions, and maximum injection time to 45 ms. No lock masses were included.

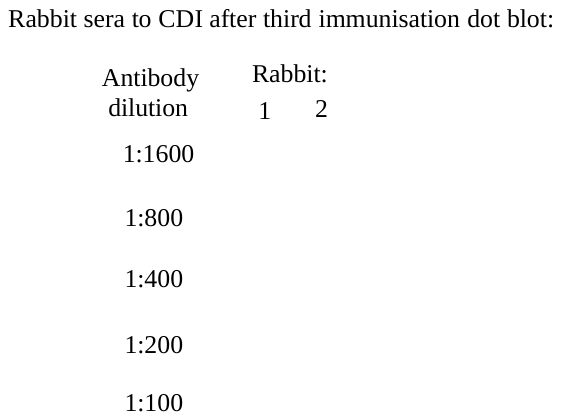

**Figure S1: Dot Blot analysis of binding of Brosi_A1236 polypeptide cys-DIREITGVASDR to rabbit sera after third immunization with polypeptide. Analysis provided by supplier (SABio, Singapore)**

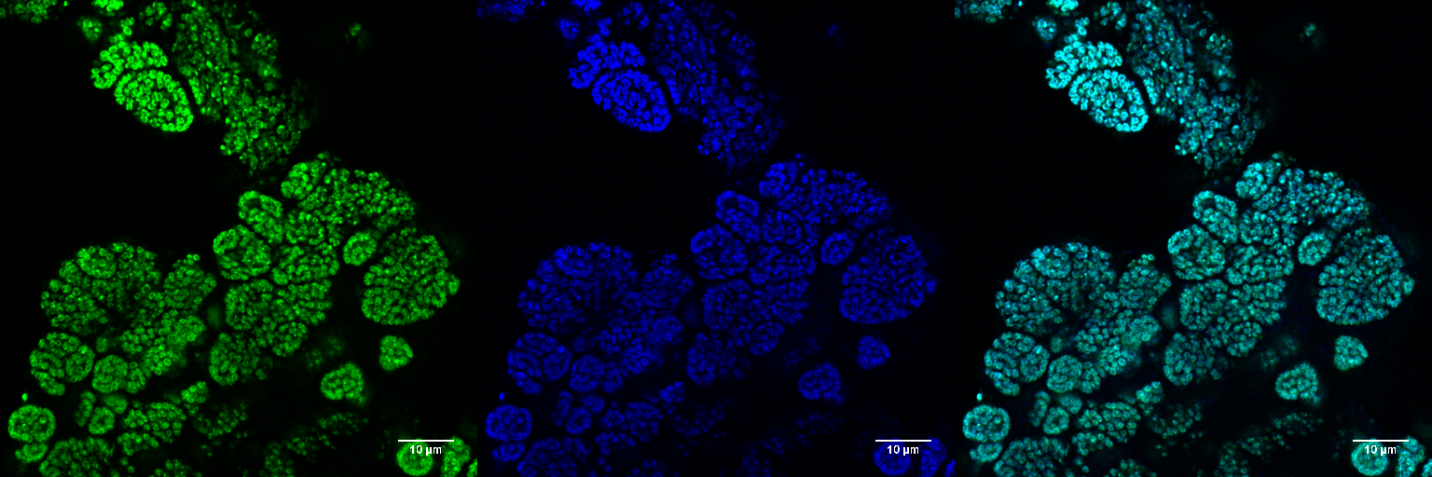

**Figure S2: Fluorescence in situ hybridization (FISH) image of the *Candidatus* Brocadia sinica-enriched Anammox biofilm with probes probe Bsi630 (green) targeting B. sinica and general Anammox probes Amx820 and Amx1900 targeting all Anammox bacteria (blue). Cyan represents overlap of green and blue showing complete dominance of B. sinica.**

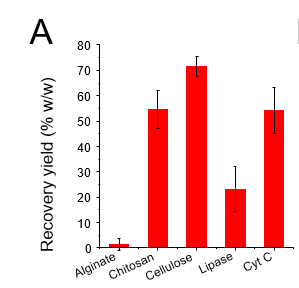

**Figure S3: Recovery yield of representative exopolymers (anionic (alginate, lipase), cationic (cytochrome C (Cyt C), chitosan) and neutral (cellulose)) following dissolution in ionic liquid (EMIM-Ac/DMAc) and recovery by ethanol precipitation.**

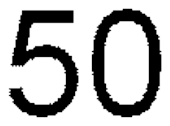

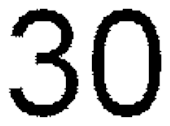

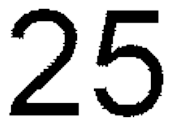

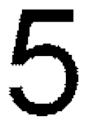

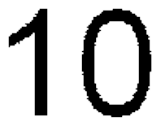

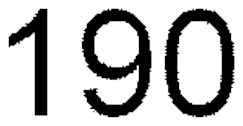

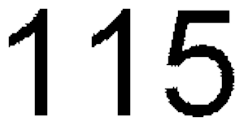

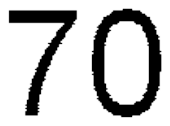

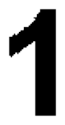

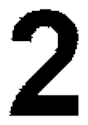

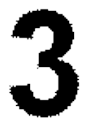

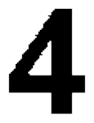

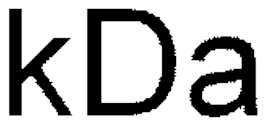

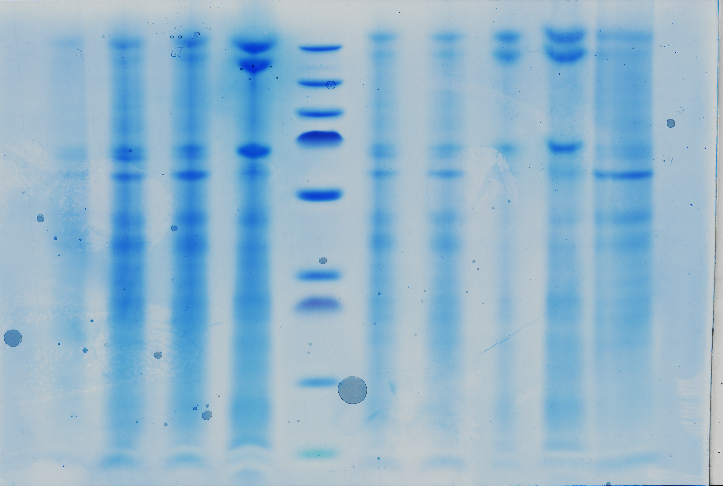

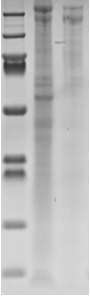

**Figure S4: SDS-PAGE gel showing extracellular proteins extracted from *Ca.* B. sinica-enriched granules using EMIM-Ac/DMAc (IL-EPS, Lane 2), anion exchange chromatography (AEC)-purified anammox protein extract (EPI, Lane 3), and similar molecular weight profile was observed in IL-GEL (Lane 4). Lane 1 is the PageRuler™ prestained protein ladder.**

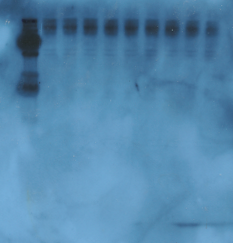

**Figure S5: Immunoblotting validation by Western blot showing positive blot of Brosi_A1236 doublet to cys-DIREITGVASDR antibody.**
